# Local translation in synaptic mitochondria influences synaptic transmission

**DOI:** 10.1101/2020.07.22.215194

**Authors:** Roya Yousefi, Eugenio F. Fornasiero, Lukas Cyganek, Stefan Jakobs, Silvio O. Rizzoli, Peter Rehling, David Pacheu-Grau

## Abstract

Mitochondria possess a small genome that codes for core subunits of the oxidative phosphorylation system, and whose expression is essential for energy production. Information on the regulation and spatial organization of mitochondrial gene expression in the cellular context has been difficult to obtain. Here we addressed this by devising an imaging approach to analyze mitochondrial translation, by following the incorporation of clickable non-canonical amino acids. We applied this method to multiple cell types, including hippocampal neurons, where we found ample evidence for mitochondrial translation in both dendrites and axons. Translation levels were surprisingly heterogeneous, were typically stronger in axons, and were independent of their distance from the cell soma, where mitochondria presumably descent from. Presynaptic mitochondrial translation correlated with local synaptic activity, and blocking mitochondria translation reduced synaptic function. Overall, these findings demonstrate that mitochondrial gene expression in neurons is intimately linked to neuronal function.

## Introduction

Mitochondria support the metabolic and bioenergetics demands of cells. The mitochondrial oxidative phosphorylation system (OXPHOS) transforms energy from reducing equivalents into ATP. During evolution, mitochondria retained a genome (mtDNA), which encodes a small set of proteins (thirteen proteins in human), all of which are core components of the OXPHOS system. All other mitochondrial proteins are nuclear-encoded, synthesized on cytosolic ribosomes, and transported into mitochondria (Wiedemann and Pfanner, 2017; Grevel et al., 2019; Hansen and Herrmann, 2019; Dennerlein et al., 2017). Accordingly, the OXPHOS system is formed from subunits of dual genetic origin. The challenge of this compound system is that cells need to adapt the availability of the nuclear- and mitochondria-encoded proteins to each other, in order to generate membrane protein complexes of defined stoichiometry. Mitochondrial gene expression requires DNA replication, transcription, and translation of the mtDNA. Approx. 25% of the mitochondrial proteome participates in these processes (Richter-Dennerlein et al., 2015; Morgenstern et al., 2017; Sickmann et al., 2003). Given the importance of gene expression in mitochondria, it is not surprising that dysfunction in any of these processes causes severe human disorders, the so-called mitochondrial disorders, which frequently affect the central nervous system (Ghezzi and Zeviani, 2018; Suomalainen and Battersby, 2018; Area-Gomez et al., 2019; DiMauro, 2019). However, tissue-specific pathologies of mitochondrial diseases are still not well understood, especially because the local regulation of these mitochondria processes is relatively unclear.

Neurons represent highly polarized cells in which the cell body, axon and dendrites display distinct morphologies and signaling functions. Mitochondria provide more than 90% of the ATP that is required for neuronal activity, and are thus essential for synaptic transmission (Harris et al., 2012). Moreover, the presence of mitochondria at synaptic boutons has been suggested to regulate the size of the synapse and the number of vesicles (Smith et al., 2016), and these mitochondria also appear to present a distinct proteome, specialized for energy supply (Völgyi et al., 2015). This finding indicates that mitochondria may undergo function-specific remodeling in order to adapt to distinct subcellular processes. Indeed, late endosomes bearing nuclear encoded mRNAs have been shown to pause at mitochondria and constitute a platform to translate mitochondrial proteins to support their function in axons (Cioni et al., 2019). Moreover, the presence of cytosolic ribosomes, transcripts, and local cytosolic protein synthesis has been demonstrated in presynaptic terminals, further supporting the idea of compartment-specific neuronal plasticity (Hafner et al., 2019; Kuzniewska et al., 2020). However, it is still unclear whether mitochondria are able to synthesize proteins in the different neuronal compartments, distant from the cell body, and whether and how this process influences synaptic activity.

To answer this question, we established an imaging technique that allows to monitor mitochondrial translation in the cellular context. This technique demonstrated that neuronal mitochondria synthesize proteins in regions distant from the cell body, albeit not all mitochondria translate to the same extent. Translation in axonal mitochondria is higher than that observed in dendritic mitochondria. Moreover, the rate of mitochondrial translation at synaptic terminals correlated with synaptic activity, and inhibiting mitochondrial translation reduced synaptic function. Altogether, our analysis shows that mitochondrial gene expression is a compartment-and function-dependent process in neurons, indicating a novel layer of neuronal activity regulation.

## Results

### Visualizing mitochondrial translation in the cellular context

The standard procedure to analyze mitochondrial translation is radiolabelling of newly synthesized polypeptides in intact cells or in purified mitochondria. However, this approach does not allow an assessment of translation in the context of the cellular anatomy, since it lacks spatial resolution. To monitor mitochondrial protein synthesis in the context of an individual cell and to obtain quantitative information on the translation process, we devised an imaging approach, based on the FlUorescent Non– Canonical Amino acid Tagging (FUNCAT) procedure (Dieterich et al., 2010). Cytosolic translation was selectively inhibited by treatment with the antibiotic harringtonine, and cells were subsequently incubated with the alkyne-containing non-canonical amino acid L-homopropargylglycine (HPG). Under these conditions, HPG is specifically incorporated into mitochondrial translation products, and can be visualized by a subsequent click reaction to azide-containing fluorescent dyes (**Fig 1A**). Using this approach, we monitored mitochondrial protein synthesis in human skin fibroblasts, and obtained HPG-specific fluorescent signals that colocalized well with mitochondria. As expected for mitochondrial translation products, the fluorescent signals were sensitive to the mitochondrial translation inhibitor chloramphenicol (**Fig 1B**). Similarly, when we monitored mitochondrial protein synthesis in cells lacking mtDNA (Rho0 cells) HPG-specific fluorescent signals were lost (**Fig 1C**), suggesting that HPG is specifically incorporated into mitochondrial translation products under these experimental conditions.

**Figure 1.**
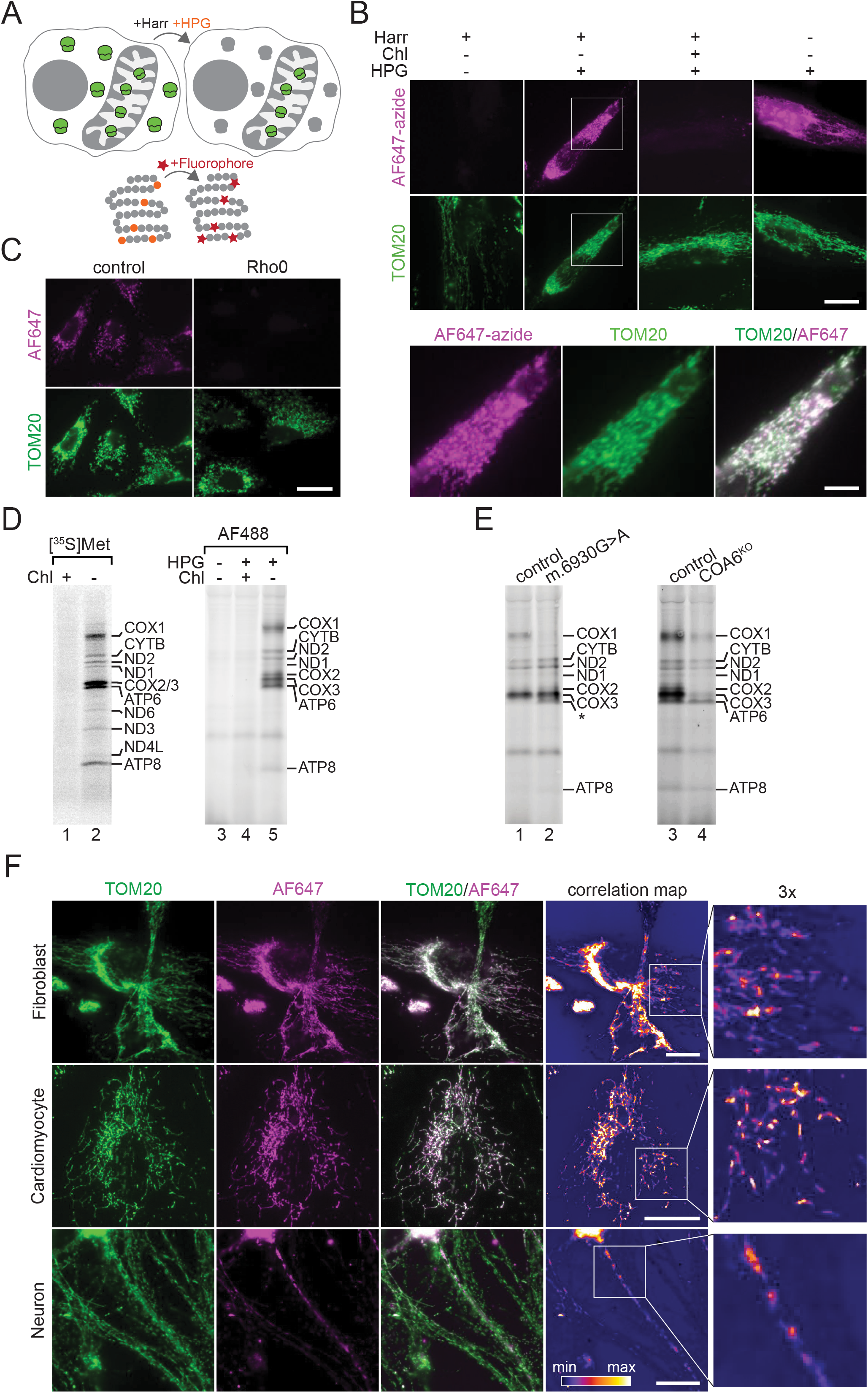
Visualizing mitochondrial translation in the cellular context. **A,** Schematic presentation of method. Cytosolic translation is inhibited with Harringtonine (Harr) while mitochondrial ribosomes are allowed to incorporate the alkyne-containing Methionine homolog (HPG, orange) in their newly-synthesized proteins. The HPG moieties are later clicked to an azide-conjugated fluorophore (star, red), through a copper-catalyzed Huisgen cycloaddition (click) reaction, and can be visualized under the microscope. **B,** Newly-synthesized proteins were labelled in WT fibroblasts with AF647-azide (magenta), using the method described in **A**. The addition of chloramphenicol (Chl) inhibits translation in the mitochondria, confirming the specificity of the click reaction. TOM20 was used as a mitochondrial marker (green). The magenta image shown in the absence of both inhibitors (the rightmost image) was acquired with a shorter exposure, due to the high signal in the cytosol under these conditions. Scale bar, 30 μm, and 10 μm for the zoom panels (bottom). **C,** Mitochondrial translation products were labeled in the presence of HPG and AF647 (magenta) in osteosarcoma 143B control and Rho0 cell lines. Scale bar, 25 μm. **D,** A comparison of mitochondrial translation labeling using either radioactive ^35^S-Met in WT HEK293T cells, or HPG followed by click reactions with AF488-azide, in isolated mitochondria. The samples were analyzed by SDS-PAGE and either revealed with digital autoradiography or fluorescent gel imaging. Both labelling methods show very similar patterns. **E,** Similar experiments, performed by click reactions in 143B COX1 mutant cells (m.6930G>A), derived from a patient, or in COA6^KO^ HEK293T cells, in comparison to the respective controls. **F,** Mitochondrial translation products were labeled in the presence of HPG and AF647 (magenta) in fibroblasts, iPSCs-derived cardiomyocytes and hippocampal neurons. TOM20 was used as a mitochondrial marker (green). Scale bar, 25 μm. The fire look-up table shows the correlation of AF647 and TOM20 channels.

To test whether these translation products conform to the expected nature of the OXPHOS subunits, we compared our results with the classical method of pulse labelling with [^35^S] methionine, followed by SDS-PAGE and autoradiography (**Fig 1D**).

HGP-labeled proteins were detected with AF488-azide, and provided a similar pattern to [^35^S] methionine labeling, which was, as expected, sensitive to chloramphenicol (**Fig 1D**). To confirm the identity of selected proteins, we first used a mutant cell line with a defect in mitochondrial translation of COX1 (m.6930G>A). In these cells the full length COX1 protein is not translated (Bruno et al., 1999; Richter-Dennerlein et al., 2016). This was correctly reported by the HPG-based analysis (**Fig 1E**, left panel). Second, it has been demonstrated that the absence of the thiol reductase COA6 strongly decreases COX2 stability, and thereby indirectly affects the abundance of COX1 and COX3 (Soma et al., 2019; Stroud et al., 2015; Pacheu-Grau et al., 2015; 2020). HPG-dependent labelling of mitochondrial translation products in COA6 knock-out cells (COA6^KO^) resulted in the expected pattern of the loss of COX2, COX1 and COX3 (**Fig 1E**, right panel). Overall, these experiments demonstrate that HPG labelling enables the specific detection of mitochondria-encoded polypeptides.

Mitochondria are optimized to fulfill the specific metabolic demands of different cell types. While it is becoming evident that mitochondrial gene expression is able to adapt to cell and tissue demands (Williams et al., 2018; Mootha et al., 2003; Di Bartolomeo et al., 2020), it is still unclear how mitochondrial gene expression is spatially coordinated to respond to differential demands within separate areas of single cells. To approach this question, we analyzed mitochondrial gene expression in three different cell types: human fibroblasts, human iPSCs-derived cardiomyocytes, and rat hippocampal neurons. Interestingly, by correlating newly synthesized translation products with the mitochondrial marker TOM20, we identified regions of high and low mitochondrial translation, reflecting compartment specific mitochondrial translation in the cellular context of the different cell types (**Fig 1F**). This effect was most striking for neurons, where profound differences could be noted between the spatial pattern of the mitochondria and that of mitochondrial translation (**Fig 1F**).

In summary, these experiments validate an imaging strategy to visualize mitochondrial translation in cells, which can be applied to determine and quantify mitochondrial gene expression in distinct cellular regions.

### Translating mitochondria distribute along neuronal branches

Most mitochondrial proteins are nuclear-encoded, synthesized on cytosolic ribosomes, and imported into mitochondria. In neurons, some nuclear-encoded mRNAs travel along axonal branches and are translated locally at their destination (Cioni et al., 2019; Turner-Bridger et al., 2018; Shigeoka et al., 2016). However, most transcripts are thought to be translated in the vicinity of the cell body. Since OXPHOS complexes are formed from mitochondrial- and nuclear-encoded subunits, we first addressed the question of whether mitochondria distant from the cell body can synthesize mtDNA-encoded proteins. Importantly, this process has not been detected in the past, even in experiments specifically analyzing local translation in neurites. It is possible that in such experiments the mitochondrial translation was simply too low to be detected, in the context of the substantially higher levels of non-mitochondrial translation (see **Fig 1B,** the non-mitochondrial translation levels are shown in the right-most panels).

To test this, we labelled mitochondrial translation products with HPG in hippocampal neurons. To define individual neurons and identify the different cellular compartments, we used the lipophilic dye DiO ((Cheng et al., 2014; Honig and Hume, 1986). Although the most intense signal for mitochondrial translation was present in proximity to the cell body, mitochondrial translation was apparent at various positions of neural branches, distant from the cell body (**Fig 2A**). Since mitochondria travel along the axon to provide energy for synaptic processes (Chang et al., 2006; Chada and Hollenbeck, 2004), we considered that the observed mitochondrial translation products could have been synthesized in the cell body, before being transported in the neurites. To test this possibility, we analyzed the mitochondrial translation levels after an HPG pulse of only 5 minutes, which does not allow for substantial transport. The overall pattern of labeling was similar to the one observed with a 30-minute pulse (**Fig 2b**), and neither showed a substantial decrease in translation with distance from the cell body (**Fig 2C**). Moreover, we also labelled mitochondrial translation products in the presence of nocodazole, an inhibitor of microtubules polymerization, to block transport of mitochondria along neuronal branches (Hoebeke et al., 1976) (Kuromi and Kidokoro, 1998).The pattern or labeling was again similar (**Fig 2B,C**), implying that mitochondrial translation can occur distant from the cell body, and is independent of transport along neurites.

**Figure 2.**
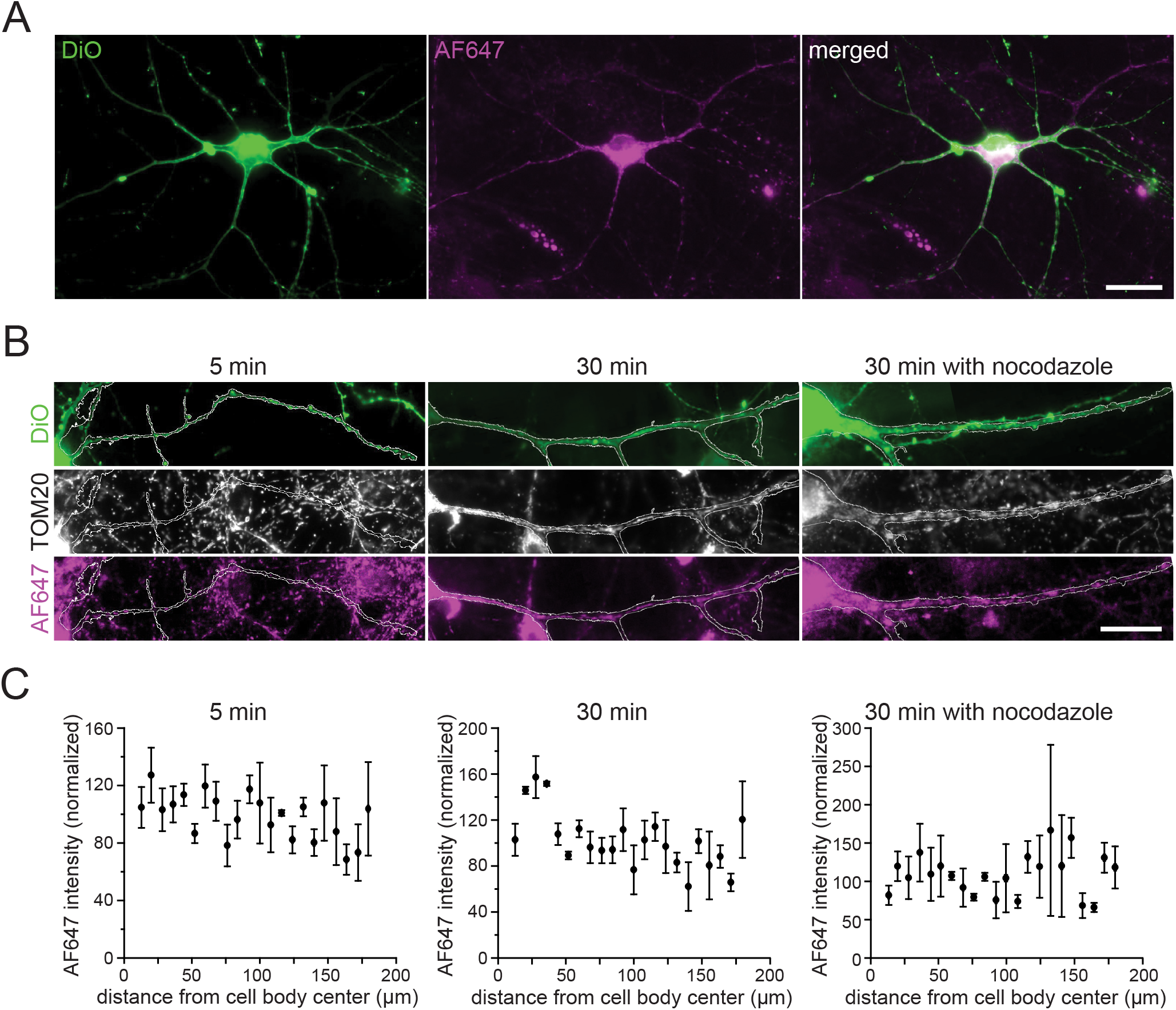
Translating mitochondria are distributed along neural branches. **A,** Mitochondrial translation was labeled using HPG and AF647 (magenta) in hippocampal neuron cultures. Individual neurons were visualized using the lipophilic fluorescent crystal dye (DiO, green). **B,** Representative images of actively translating mitochondria along the neurites after labeling with HPG and AF647 for 5 min, 30 min and in the presence of nocodazole. TOM20 was used as a mitochondrial marker (grey). The outline of one neurite is included in all channels. Scale bar, 25 μm. **C,** Quantification of mitochondrial translation signals in relation to the distance from the cell body, in the same conditions as in panel B (Mean ± SEM, normalized to the median of all data points, N=3 independent experiments, with 10 images per experiment).

### Axonal mitochondria display increased translation

As the nature of the neurites could not be assessed in the previous experiments, we performed a more specific assay to differentiate them, based on immunostaining the samples for ANK-G, a protein located in the axon initial segment. This approach enabled us to differentiate between axons and dendrites, as neurites containing ANK-G, or devoid of it. Initial segments of the neurites, protruded immediately from the soma, were then imaged (**Fig 3A**). An analysis of the mitochondria demonstrated that the axonal ones translate more than the dendritic ones (**Fig 3B**), thus suggesting that the levels of mitochondrial translation depend on the nature of the cellular compartment containing them.

**Figure 3.**
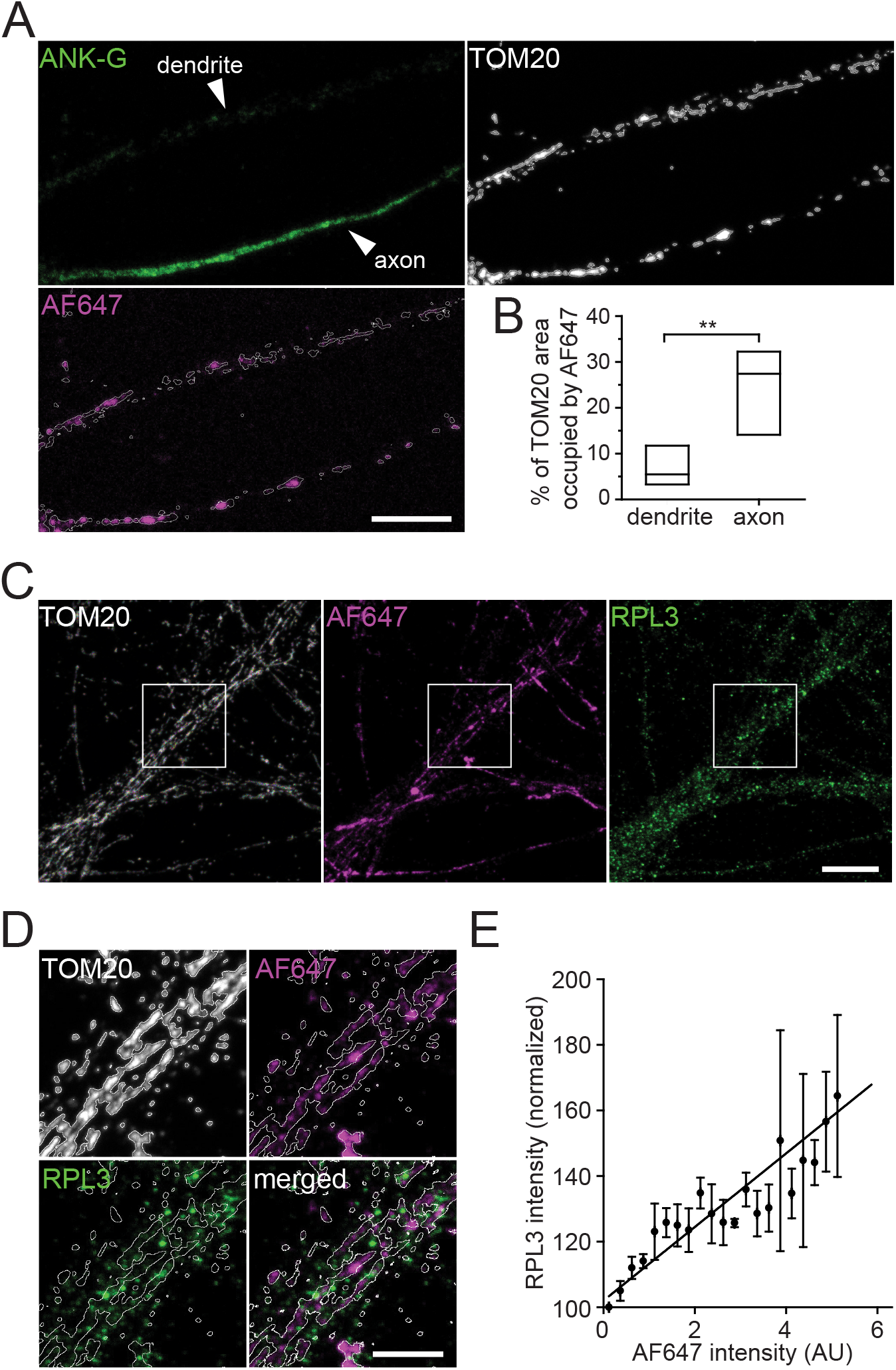
Axonal mitochondria displayed increased translation. **A,** Representative confocal images of mitochondrial translation using HPG and AF647 in hippocampal neurons (magenta). TOM20 was used as a mitochondrial marker, and ANK-G was used as an axonal marker. Scale bar, 10 μm. **B,** Quantification of the percentage of mitochondria occupied by translation products in axons and dendrites (N=5 independent experiments, with 10 images per experiment; *p<0.01, t-test). **C,** Representative confocal images of mitochondrial translation using HPG and AF647 in hippocampal neurons (magenta). RPL3 (green) was used to visualize cytosolic ribosomes and TOM20 (grey) was used as a mitochondrial marker. Scale bar, 10 μm. **D,** A closer view of the selected square in panel A, showing cytosolic ribosomal particles (RPL3) in association with actively translating mitochondria (AF647). TOM20 was used to identify the mitochondria. Scale bar, 5 μm. **E,** Correlation analyses of mitochondrial translation levels with the amount of RPL3 bound to mitochondrial particles (Mean ± SEM, normalized to the lowest value, N=3 independent experiments, with 15 images per experiment, R^2^= 0.8 and *p<0.0001).

Mitochondrial translation is regulated by the influx of nuclear encoded OXPHOS subunits (Richter-Dennerlein et al., 2016; Couvillion et al., 2016). Indeed, cytosolic ribosomes have been found associated with mitochondria, suggesting that translation of selected nuclear-encoded proteins may occur in proximity to the mitochondrial surface (Gold et al., 2017; Suissa and Schatz, 1982; Costa et al., 2018; Williams et al., 2014). As our analyses demonstrated that different neuronal areas contained mitochondria with higher or lower translation signals (**Fig 1F**), we expected that the levels of attached cytosolic ribosomes may also vary. To test this, we labelled mitochondrial translation products as above, and used TOM20 and RPL3 as markers for mitochondria and cytosolic ribosomes, respectively. Cytosolic ribosomes could be visualized in proximity to newly synthesized mitochondrial translation products in neuronal branches (**Fig 3C, D**), and their amounts correlated to the translation levels (**Fig 3E**). These data suggest that the translation of both mitochondrial- and nuclear-encoded mitochondrial proteins can be achieved locally at mitochondria in the neurites.

### Presynaptic activity correlates with protein synthesis in mitochondria

Presynaptic cytosolic translation plays a role in axonal development and synaptic plasticity (Schanzenbächer et al., 2018; Hafner et al., 2019; Yoon et al., 2012; Hillefors et al., 2007). However, it is still unclear whether the mitochondria located in the presynaptic boutons also translate actively, and whether this connects to synaptic transmission. We therefore analyzed presynaptic boutons, taking advantage of HPG labeling in combination with STED microscopy. As a marker for presynaptic sites, we labeled recycling vesicles using an antibody for the lumenal (intravesicular) domain of the vesicle marker synaptotagmin 1 (SYT1), conjugated to the fluorophore Atto590 (Truckenbrodt et al., 2018). The SYT1 epitope is exposed by the actively recycling vesicles during exo- and endocytosis, and enables the internalization of the antibody. The neurites were stained with DiO, and were classified as axonal or dendritic based on their morphology (the presence or absence of dendritic spines) and based on the presence of SYT1-positive vesicle clusters within the DiO-marked volumes. Mitochondrial translation was readily detected (**Fig 4A**).

**Figure 4.**
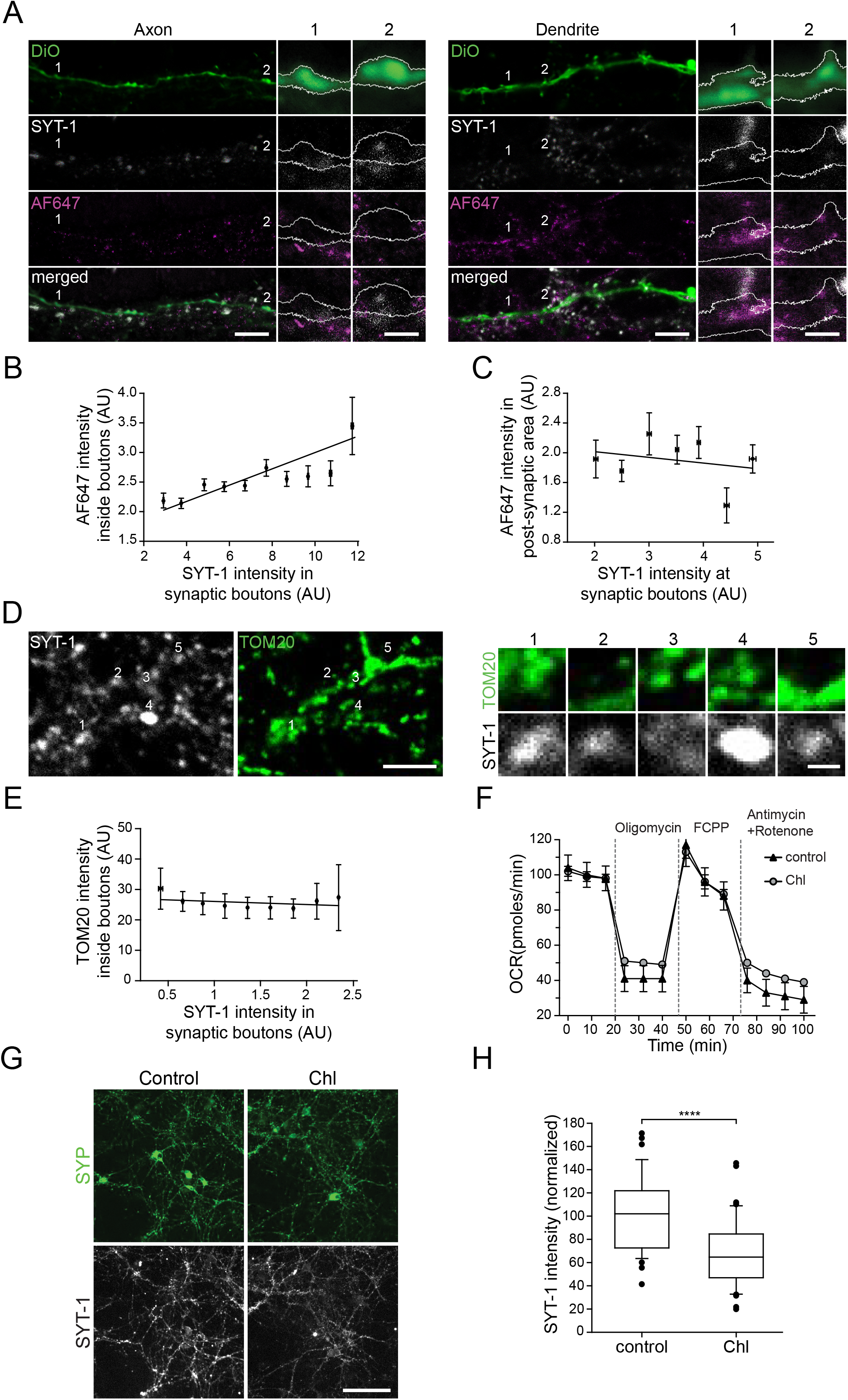
Protein synthesis in mitochondria correlates with presynaptic activity. **A,** Representative images of axons (left) and dendrites (right) in which mitochondrial protein synthesis is labeled with HPG and AF647 (magenta, STED). Active synaptic boutons are labeled by the uptake of SYT1-Atto590 (grey, STED), which reveals actively recycling vesicles. The plasma membrane is labeled using DiO (green). Scale bar, 5 μm. Two regions of interest from each neurite are zoomed in. The white lines indicate the DiO outline. Scale bar, 1 μm. **B-C,** Correlation analyses of mitochondrial translation levels with SYT1 in synaptic boutons (panel **B**; R^2^= 0.78 and *p<0.01) and in adjacent dendritic regions (panel **C**; R^2^= 0.063 and p>0.05; N=3 independent experiments, with 20 axons and dendrites per experiment)). **D,** An overview of SYT1 uptake in relation to mitochondria positions. There is no obvious correlation between the strength of the SYT1 label, indicating synaptic activity (gray), and the strength of the mitochondria staining (green), as indicated also by the five high-zoom examples. Scale bar, 1μm. **E,** Analysis of the relation between the mitochondrial staining (TOM20) and the SYT1 signals. No significant correlation could be found; p>0.05; N=5 independent experiments). **F,** Respiratory chain activity was assessed by Real-Time respirometry (Seahorse measurement) in hippocampal neurons incubated in the presence or absence of chloramphenicol for one hour (Mean, SEM, N = 6). OCR, oxygen consumption rate. **G,** Representative images of hippocampal neurons incubated in the presence or absence of Chloramphenicol (Chl) for 1 hour. Live neurons were then labeled with SYT1-Atto647N antibodies (SYT1, grey) for 10 min in Tyrode’s buffer. Following fixation, synaptophysin (SYP) was used as a presynapse marker (green). Scale bar, 100 μm. **H,** Quantification of SYT1 intensity in control and Chloramphenicol-treated samples. (N = 3 independent experiments, with 20 images per experiment). The values in each experiment were normalized to the mean of the respective control experiment. The difference was significant. *p<0.0001, Mann-Whitney U tests).

As oxidative phosphorylation is required to provide energy for synaptic activity (Sun et al., 2013; Ivannikov et al., 2013), it is probable that mitochondrial biogenesis and the production of OXPHOS subunits need to scale with the levels of synaptic activity. Since the SYT1 staining provides a simple estimate of the local levels of activity, we compared the intensity of this staining to the local mitochondrial translation levels. The two correlated significantly (**Fig 4b**). No correlation could be found between the synaptic activity levels and translation in the dendritic segments closely apposed to the respective presynapses (**Fig 4c**), suggesting that presynaptic activity is specifically correlated to local protein production in the presynapse. This correlation does not simply reflect a link between the presence of mitochondria and synaptic activity, since presynaptic boutons can have widely varying levels of both activity and mitochondria (**Fig 4D**), and since no correlation could be found between the two (**Fig 4E**).

To test the strength of this interrelation, we inhibited mitochondria translation using chloramphenicol. A chloramphenicol treatment of neurons did not affect mitochondrial respiratory chain function (**Fig 4F**). We then monitored presynaptic activity using the SYT1 assay upon chloramphenicol treatment (**Fig 4G**). This inhibition resulted in a significant reduction of presynaptic activity, suggesting that mitochondria translation is strongly linked to neuronal function (**Fig 4H**).

## Discussion

We developed here an imaging approach, based on the FUNCAT procedure (Dieterich et al., 2010) which enabled us to identify specifically the production of proteins encoded by the mitochondrial genome. When applied to fibroblasts or iPSC-derived cardiomyocytes, our approach demonstrated high levels of mitochondrial translation, which correlated well with the mitochondrial size and density (**Fig 1F**). In neurons, however, a far more heterogeneous image was obtained (**Fig 1F**), with some mitochondria being substantially more active than the rest, especially in axons.

The consensus view is that mitochondria typically “descend” from the cell body, and are then trafficked to different dendritic and axonal sites (Devine and Kittler, 2018). Substantial research has been conducted on the mechanisms of mitochondria trafficking, resulting in substantial knowledge on both the motors involved and on the molecules coordinating mitochondrial traffic, as the mitochondrial Rho GTPases (MIROs)(Devine et al., 2016). Overall, a view emerges of mitochondria as dynamic organelles that can be trafficked at speeds of μm per second, and whose mobility is directed by the neuronal activity (Sajic et al., 2013). As mitochondrial proteins are among the most long-lived in the brain (Fornasiero et al., 2018), this view could easily be reconciled with a concept of mitochondrial proliferation in the cell body, followed by usage in neurites, for long time periods, without a need for local mitochondria protein production and turnover.

This view agrees with much of the literature, including the observation that local protein synthesis in pre- and postsynaptic compartments appeared to be resistant to chloramphenicol (Hafner et al., 2019), and was therefore independent of mitochondrial translation. However, a detailed measurement of mitochondria motility during neuronal maturation suggested that mitochondria become less mobile as the neurons age, and that they tend to become localized at synapses (Lewis et al., 2016). Such mitochondria would eventually need to replace their proteomes locally, which induces a need for both local translation of proteins on the basis of the nuclear genome (followed by mitochondrial import), and local translation of proteins encoded by the mitochondrial genome. We found evidence for both processes, and we suggest them to be quantitatively correlated (**Fig 3E**). This would be expected for the formation of functional mitochondria, which need substantial levels of both nucleus- and mitochondria-encoded elements. Some previous evidence supporting this conclusion could be found in transcriptomics and proteomics analyses of synapse-enriched fractions, which indicated that nucleus-encoded transcripts for mitochondria proteins are enriched in synapses(Hafner et al., 2019), where they may be translated (Kuzniewska et al., 2020)

The functional implications of local mitochondrial translation are still difficult to assess, beyond the simple replacement of old and potentially damaged components. Mitochondria in synapses participate not only in energy production, but also in calcium buffering, implying that they take an active role in neuronal function and dysfunction (Frere and Slutsky, 2018). This implies that mitochondria in or near active synapses, especially in the presynaptic compartment, would be more often required to engage in their buffering function in active terminals. It is probable that an organelle or protein complex that is required to function more frequently, and undergoes repeated changes in its conformation, has a higher chance of becoming damaged, and therefore of needing to be replaced by newly synthesized proteins. This would explain the link between synaptic activity and presynaptic production of mitochondrial proteins (**Fig 4**), albeit we would like to point out that there is no strong link yet between protein usage and protein damage or replacement in mitochondria (although previous work has demonstrated such a phenomenon for presynaptic vesicles (Truckenbrodt et al., 2018). Overall, our work provides a first view of mitochondrial translation in complex cells, demonstrating that this process can take place far from the cell body, and can be tuned in relation to location (*e.g.* dendritic or axonal) and to cellular activity.

## Material and Methods

### Cell lines and cell culture

All non-primary cell lines were cultured in Dulbecco’s modified Eagle’s medium (DMEM, ThermoFischer) and were supplemented with 10% (V/V) heat-inactivated fetal bovine serum (Biochrom, Berlin, Germany), 2 mM L-glutamine, 1 mM sodium pyruvate and 50 μg/ml uridine. They were incubated at 37°C with 5% CO_2_ and passaged on a regular basis. HEK293T Flp-In™ T-REX™ were purchased from Invitrogen. WT fibroblasts, COA6^KO^ HEK293T, WT and Rho0-143B cell lines were previously reported (Pacheu-Grau et al., 2018; 2020; Gómez-Durán et al., 2010; King and Attardi, 1989). The G6930A 143B cell line was a gift from G. Manfredi and J. Montoya (Bruno et al., 1999; Richter-Dennerlein et al., 2016). For imaging purposes, the cells were seeded on PLL-coated glass coverslips (0.1 mg/ml) one day before the start of the experiments.

The use of human iPSC lines was approved by the Ethics Committee of University Medical Center Göttingen (approval number: 10/9/15) and was carried out in accordance with the approved guidelines. Written informed consent was obtained from all participants prior to the participation in the study.

Human iPSC lines from a healthy donor (isWT11.8/UMGi130-A) were used in this study. Human iPSCs were maintained on Matrigel-coated plates (growth factor reduced, BD Biosciences), passaged every 4-6 days with Versene solution (Thermo Fisher Scientific) and cultured in StemMACS iPS-Brew XF medium (Miltenyi Biotech) supplemented with 2 μM Thiazovivin (Merck Millipore), for 24 h after passaging with daily medium change. Directed differentiation of human iPSCs into iPSC-cardiomyocytes was performed via WNT signaling modulation and subsequent metabolic selection, as previously described (Cyganek et al., 2018). Differentiated iPSC-cardiomyocytes were cultured for two months and were subjected to molecular and functional analyses. Cell cultures were incubated in a humidified incubator with 5% CO_2_ at 37°C.

Neuronal hippocampal cultures were obtained from dissociated hippocampi of new-born rats, as previously published (Banker and Cowan, 1977; Kaech and Banker, 2006; Truckenbrodt et al., 2018). For imaging purposes, the cells were plated at a concentration of ~30,000/cm^2^ on coverslips coated with 1mg/ml PLL, after being treated as previously described (Truckenbrodt et al., 2018). Following dissection and plating, the cells were left to adhere for 1-4 h at 37°C in a cell incubator, with 5% CO_2_. After adhesion, the medium was replaced to Neurobasal-A medium (Gibco, Life Technologies, Carlsbad, CA, USA), containing 1:50 B27 supplement (Gibco) and 1:100 GlutaMAX (Gibco). Neurons were kept in culture at 5% CO2 and 37°C, for 14-21 days before use.

### *In vivo* labeling of newly synthesized mitochondrial-encoded peptides with HPG

To label newly synthesized mitochondrial-encoded peptides, a click chemistry-based approach was adapted (Estell et al., 2017). Briefly, cells were transferred to methionine-free medium. In neurons, the labeling was done in warm Hank’s Balanced Salt Solution (HBSS, ThermoFisher). Cytosolic translation was stopped using harringtonine (200μM, Carbosynth) for 20 minutes. For control experiments blocking mitochondrial translation, cells were incubated in the presence of 75 μg/ml Chloramphenicol. 0.5 mM of L-Homopropargylglycine (HPG, ThermoFisher) was then added for 5-30 minutes. Cells were then transferred to buffer A containing 10mM HEPES, 10mM NaCl, 5mM KCl, 300mM sucrose, and 0.015% digitonin, for 2 min on ice, followed by 15 seconds in buffer A without digitonin. Coverslips were fixed using 4% PFA in PBS (137 mM NaCl, 2.7 mM KCl, 10 mM Na_2_HPO_4_, 2 mM KH_2_PO_4_) at pH 7.5 for 30 minutes at room temperature and were then further processed for click-chemistry and immunostaining. For experiments interfering with microtubule polymerization neurons were incubated with nocodazole (30 μM, for 2h).

### Click-chemistry and immunostaining of fixed cells

After formaldehyde fixation, coverslips were washed for 5 min with PBS quenched for 15 min with 100mM NH_4_Cl in PBS. Blocking and permeabilization were performed in staining solution (PBS + 5% BSA + 5% tryptone peptone + 0.1% Triton X-100), using 3 solution exchanges, each for 5 min. After a brief wash with 3% BSA in PBS, coverslips were clicked for 20 minutes using a commercial kit (Click-iT Cell Reaction Buffer Kit; ThermoFisher), with 3 mM AlexaFluor 647-azide (ThermoFisher). After another quick wash with 3% BSA in PBS, primary and secondary antibodies were diluted in the staining solution, and were applied sequentially for 1 h. Three 5-min washing steps with staining solution were performed between the primary and secondary antibody incubations. 3 x 5min sequential washes with blocking solution (PBS + 5% BSA + 5% tryptone peptone), high-salt PBS (PBS + 500 mM NaCl), and PBS were performed at the end of the staining procedure, to remove thoroughly any non-specific binding of the antibodies. The coverslips were then embedded in Mowiol (Calbiochem, Billerica, MA, USA) and were allowed to dry overnight at room temperature, before imaging.

The antibodies used in this study are as follows: TOM20 (rabbit, Proteintech), ANK-G (mouse, Antibodies incorporated), SYP (guinea pig, Synaptic systems), RPL3 (mouse, Proteintech), goat anti-Rabbit Alexa Fluor Plus 488 (ThermoFisher), donkey anti-Mouse conjugated to Cy3 (Dianova), goat anti-mouse STAR 580 (Abberior), goat anti-Guinea Pig Alexa Fluor 546 (ThermoFisher).

### Labeling of mitochondrial-encoded peptides with [^35^S]-methionine

Radioactive labeling of mitochondrial translation products was performed according to a previously published protocol (Chomyn, 1996) with minor modifications. Cytosolic translation was inhibited with harringtonine (200 μM) for 20 min. Then, 200 μCi/ml [^35^S]-methionine was added to the cells and was incubated at 37°C for 1 h. The cell lysate was loaded on a 10-18% Tris-Tricin gradient gel, and the radioactive signal was detected using Phosphor Screens and a Storm 820 scanner (GE Healthcare).

### Click-chemistry on extracted mitochondria

Cells were incubated in methionine-free medium with 200 μM harringtonine for 20 minutes, and 500 μM HPG was added to the cells for 4 hours. After harvesting the cells, mitochondria were isolated by differential centrifugation, as previously described (Lazarou et al., 2009). The protein concentration was measured using a standard Bradford assay, and 100 μg of mitochondria proteins were pelleted and dissolved in 25 μl 50mM Tris (pH 8.8) containing 1 mM PMSF and 1% SDS. A click reaction was performed on ice using a commercial kit (Click-iT Cell Reaction Buffer Kit; ThermoFisher), with 80 μM Alexa Fluor 488-azide (Sigma). According to the manufacturer’s protocol, proteins were purified from the mixture using a MeOH/chloroform approach, after the end of the click reaction. The extracted pellet was completely dried by heating up to 50°C and was then dissolved in loading buffer containing 8 M Urea, 100 mM Dithiothreitol (DTT) and 1% benzonase. It was then incubated at 37 °C for 20 minutes and was then loaded on a 10-18% Tris-Tricine gradient gel. Fluorescent signals in the gels were analyzed using a FLA-9000 Starion image scanner (Fujifilm).

### Membrane staining with DiO for visualizing single neurons

Membrane staining was performed on fixed coverslips, after click-chemistry and immunostaining, without Triton X-100 in the solutions. DiO crystals (ThermoFisher) were dissolved in dimethylformamide (DMF) to a stock concentration of 2 mg/ml, by mixing for 10 min at 50° C. The stock was kept at 4° C, in the dark, until use. A working concentration of 1 μg/ml was prepared in PBS, and was thoroughly mixed and sonicated. It was applied to the coverslips for 20 minutes at 37° C. The coverslips were washed with PBS briefly, and were left overnight in PBS at room temperature. The next day, the coverslips were embedded in Mowiol, as above, and were dried overnight before imaging.

### Live-cell immunostaining of recycling synaptic vesicles

Recycling synaptic vesicles were labelled using a mouse monoclonal antibody against the lumenal (intravesicular) domain of Synaptotagmin 1 (clone 604.2; Synaptic Systems, Göttingen, Germany) conjugated to either Sulfo-Cyanine 3 (#105 311C3) for normal epifluorescent imaging, or Atto 590 (custom made by Synaptic Systems) for STED imaging. Neurons were incubated with 8.3 μg/ml antibody in their own medium for 1 h (unless indicated differently) and were subsequently processed for metabolic labeling of new protein synthesis. For experiments analyzing effects of mitochondrial translation inhibition, neurons were incubated for 1 h in the presence of 75 μg/ml Chloramphenicol in Tyrode’s solution (124 mM NaCl, 5 mM KCl, 30 mM glucose, 25 mM HEPES, 2 mM CaCl2, 1 mM MgCl2, pH 7.4).

### Real-time respirometry

Oxygen consumption rate (OCR) was measured with a XF96 Extracellular Flux Analyzer (Seahorse Bioscience, Billerica, MA, USA) as previously described (Pacheu-Grau et al., 2020). Briefly, hippocampal neurons were seeded on a PLA-coated plate at a density of 15.000 cells/well and grown in the plate for 16 days. Mitochondrial translation was inhibited by incubating the cells with 75 μg/ml Chloramphenicol for 1 hour before starting the experiment. Baseline respiration was measured in neuronal media after calibration at 37 °C in an incubator without CO_2_. Periodic measurements of oxygen consumption were performed and OCR was calculated from the slope of change in oxygen concentration over time. Metabolic states measurements were performed after subsequent addition of 3 μM oligomycin, 6 μM carbonyl cyanide 4 (trifluoromethoxy)phenylhydrazone (FCCP), 1 μM antimycin A, and 2 μM rotenone.

### Image acquisition

Large field images were taken with an inverted Nikon Ti epifluorescence microscope (Nikon Corporation, Chiyoda, Tokyo, Japan) equipped with a Plan Apochromat 60×, 1.4 NA oil immersion objective, an HBO-100W Lamp, and an IXON X3897 Andor (Belfast, Northern Ireland, UK) camera, operated via the NIS-Elements AR software (version 4.20; Nikon).

The confocal images of Fig 3A were taken by a TCS SP5 STED microscope (Leica, Wetzlar, Germany) equipped with a HCX Plan Apochromat 100×, 1.4 NA oil STED objective, and operated with the LAS AF imaging software (version 2.7.3.9723; Leica). For the confocal images of Fig 3c, an Abberior microscope operated with Imspector imaging software (Abberior Instruments, Göttingen, Germany) was used. This setup was built on an Olympus IX83 base, equipped with a UPlanSApo 100× oil immersion objective (Olympus Corporation, Shinjuku, Tokyo, Japan) and an EMCCD iXon Ultra camera (Andor, Belfast, Northern Ireland, UK). Pulsed 561-nm and 640-nm lasers were used for excitation of Atto590 and Alexa Fluor 647. For high resolution images in Fig 4, the STED setup on the Abberior microscope, with easy3D-module lasers at 595 and 775 nm for depletion, was used.

### Image analysis

To analyze the distribution of translating mitochondria along the neural processes in Fig 2, the center of the soma in the images were manually marked. Using a self-written Matlab script, in which the neurite regions were selected by applying an empirically-determined threshold in the DiO channel. Mitochondrial particles within the neurite regions were selected based on the TOM20 staining, again using an empirically-determined threshold. The distance from the soma and the amount of AF647 signal in each mitochondrion were then obtained and plotted.

To analyze the mitochondrial translation in axonal and dendritic branches in Fig 3b, we used a self-written Matlab script. Mitochondria were identified, as above, by applying an empirically-determined threshold in the TOM20 channel. The ANK-G signal in the different mitochondria-marked regions was used to differentiate between axonal and dendritic mitochondria. The AF647 signal was then measured inside each identified mitochondrion.

In Fig 3e a self-written Matlab script was used to analyze the correlation of mitochondrial translation and the binding of mitochondrial ribosomes to the mitochondria. Mitochondrial particles were selected using thresholding in the TOM20 channel, as above. The signals in RPL3 and AF647 channels were measured for each mitochondrion and were plotted against each other.

To analyze the correlation between synaptic activity and mitochondrial translation in Fig 4b and c, we used a self-written Matlab script, in which we analyzed the signal in synaptic boutons, as revealed by the Synaptotagmin 1 staining, which was subjected to an empirically-defined threshold. The immediate surrounding area covered by DiO (from axonal DiO staining) was added to the bouton area, in order to analyze axonal regions. Alternatively, for dendritic regions we considered Synaptotagmin 1-defined boutons, and then identified the DiO-stained neurites that were in their immediate vicinity. In both cases, the intensities of AF647 and Synaptotagmin 1 were obtained and were plotted against each other. In Fig 4h, a self-written Matlab script was used to analyze the synaptic activity, in which the synaptic boutons were selected by the Synaptophysin signal, after being subjected to an empirically-defined threshold. The Synaptotagmin 1 signal was accordingly measured in defined areas of control and chloramphenicol-treated samples

## Statistical analysis

Two-sided Student’s t-tests or Mann-Whitney U tests were used to calculate statistical significance, as mentioned in the figure legends. A P-value of < 0.05 was considered statistically significant. The same P value was considered significant for linear regression analyses.

## Acknowledgements

Funded by the European Research Council (ERC) Advanced Grant (ERCAdG No. 339580) to PR, the SFB1286 (projects A03 (SOR), A05 (SJ), A06 (PR)) the SFB1002 S01 (LC) and CY 90/1-1 (LC), supported by the Deutsche Forschungsgemeinschaft (DFG, German Research Foundation) under Germany’s Excellence Strategy - EXC 2067/1-390729940, the German Federal Ministry of Education and Research (BMBF) / DZHK (LC), the Max Planck Society (PR, SJ) and the PhD program Molecular Biology – International Max Planck Research School and the Göttingen Graduate School for Neurosciences and Molecular Biosciences (GGNB; DFG grant GSC 226/1) (RY).

## Author contributions

RY performed the experiments, analyzed and interpreted the data and edited the manuscript. EFF provided technical advice and edited the manuscript. SJ participated in conceptualization of the work and edited the manuscript. SOR participated in data analysis and interpretation, design and conceptualization of the work and wrote and edited the manuscript. PR participated in design and conceptualization of the work, wrote and edited the manuscript and was involved in supervision during the project. DPG performed experiments, analyzed and interpreted the data, was involved in supervision during the project and wrote and edited the manuscript.

